# ABCDscores: An R package for computing summary scores in the ABCD Study®

**DOI:** 10.1101/2025.09.04.674066

**Authors:** Le Zhang, Olivier Celhay, Biplabendu Das, Samantha Berman, Laura R. Ziemer, Calen J. Smith, Anders M. Dale, Janosch Linkersdörfer

## Abstract

The ABCD Study® is the largest longitudinal research study of brain development and child health across the United States. The study generates a rich multimodal dataset, including file-based raw and derivative data and a curated tabulated dataset comprising data from a broad spectrum of research domains. Historically, summary scores in the tabulated dataset were computed using the electronic data capture system REDCap or external scripting solutions. However, this approach was rigid, unreliable, and lacked transparency around how scores were computed. Here, we introduce ABCDscores, an R software package that is used by the study to compute most non-proprietary summary scores included in the tabulated ABCD data resource and allows for customization to meet specific research needs. ABCDscores provides a flexible, open-source solution that uses R for transparent and reproducible scoring. The package is built to apply standard scoring methods across different types of assessments, ensuring consistent and accurate results even with large datasets. By simplifying complex scoring procedures into a user-friendly process, ABCDscores streamlines research workflows, reduces errors, and saves researchers valuable time. The package aims to improve analyses of data from the ABCD Study® and can serve as a model for the development of similar tools and a standardized framework for other research studies, fostering new discoveries and innovation in the field.

## Introduction

The Adolescent Brain Cognitive Development (ABCD) Study is the largest long-term study of brain development and child health in the United States, following more than 11,000 youth from ages 9–10 into early adulthood. This comprehensive, longitudinal cohort study collects a vast array of data, including neuroimaging, genetics, neurocognitive assessments, behavioral surveys, biospecimens, wearables, and environmental measures, across 21 research sites nationwide (Feldstein Ewing and Luciana 2018). Large-scale research initiatives, such as the UK Biobank and the Human Connectome Project (Sudlow et al. 2015; Van Essen et al. 2013), have established benchmarks in open science by offering comprehensive, de-identified datasets to the international scientific community, thereby expediting advancements in the field. Following this open-science model, the ABCD Study® provides its extensive, longitudinal datasets to researchers globally and promotes diverse investigations into adolescent development. The ABCD Study’s data is publicly available through the NIH Brain Development Cohorts (NBDC) Data Hub (ABCD consortium 2025).

A critical aspect of preparing ABCD data releases involves the computation of summary scores. These are composite measures derived from various assessments within the tabulated dataset. Summary scores are essential for capturing complex constructs such as cognitive functioning, mental health status, or substance use. Calculating these scores requires meticulous data processing, including handling missing data, applying appropriate scoring algorithms, and ensuring consistency across different data releases.

Originally, the ABCD Study® employed an inconsistent approach to compute summary scores. Some scores were generated as calculated fields within the Research Electronic Data Capture (REDCap) system used for data acquisition, while others were computed through external workflows utilizing various programming languages and methods. Despite its widespread acceptance in clinical research (Patridge and Bardyn 2018; Harris et al. 2009), a notable limitation of REDCap is its relatively rigid structure, which can hinder the implementation of complex algorithms. Although REDCap serves as a valuable tool for data collection and management, its built-in functionalities often fall short when researchers need to integrate or apply sophisticated computational models (Ndlovu et al. 2024). Consequently, previous solutions for computing summary scores in ABCD required the inclusion of external workflows, which, over time, resulted in inconsistent methodologies and fragmented scripts across different programming languages.

Additionally, the summary scores included in prior ABCD releases exhibited a lack of transparency and reproducibility, as the scoring algorithms were not publicly accessible and were inadequately documented. This made it challenging for data users to reproduce the scores without soliciting disclosure from the ABCD Consortium. Even if they managed to acquire the scripts, the algorithms varied between releases. In contrast to typical extensions (packages) available in popular programming languages like R or Python—distributed as versioned releases allowing users to download them and access the source code through version control platforms such as GitHub—the previous ABCD summary score scripts were poorly organized and lacked systematic version control. Together, these limitations posed significant challenges for developers aiming to extend existing algorithms and incorporate new measurements, as well as for researchers striving to reproduce the scores.

To address these challenges and streamline the research process, we introduce ABCDscores, an R package designed to automate the computation of all non-proprietary summary scores published by the ABCD Study®. In this manuscript, we detail the development and functionalities of the ABCDscores R package. We provide an overview of the included summary scores, the modular utility functions we use to compute the summaries, and discuss the implementation of standardized scoring procedures. We further provide use cases to demonstrate the package’s utility. ABCDscores aims to enhance reproducibility, allow flexibility, reduce errors, and save valuable time for researchers by encapsulating the scoring algorithms and data processing steps into a user-friendly, open-access package. We hope this manuscript will contribute to existing and future research, provide a roadmap for standardized summary score computations, and foster further innovation in this critical area.

## Methods & Implementation

ABCDscores was developed as an R package. The package is designed to automate the computation of standardized summary scores from the ABCD Study® datasets. It allows users to reproduce the summary scores that are being released as part of the ABCD tabulated data resource or to produce customized scores.

### Coding Style

ABCDscores adheres to the Tidyverse style guide for writing R code, ensuring consistency, readability, and ease of use (Wickham et al. 2019). By following this guide, we ensure that our code is not only efficient but also easily understandable, facilitating collaboration and future development.

### Core Design

The package architecture follows modular design principles, separating core functionality into distinct components to improve clarity, flexibility, and maintainability. Each summary score algorithm is implemented as a standalone function with consistent input/output patterns, making it easy for researchers to inspect the logic for a given score and incorporate specific measures into their workflows.

All scores are organized by their ABCD research domain. Currently, scores are available for the following seven domains: ABCD General (*ab*), Family, Friends & Community (*fc*), Mental Health (*mh*), Neurocognition (*nc*), Novel Technologies (*nt*), Physical Health (*ph*), and Substance Use (*su*). Within each domain, scores are organized by database table. Each table contains data for a specific survey, questionnaire, or measure administered to either the youth participant or their parent. Within each table, scores may be grouped into scales and/or subscales. For instance, the Family Environment Scale [Parent] and [Youth] tables, (*fc_p_fes*) and (*fc_y_fes*), respectively, are part of the *fc* domain. These scales include subscales like the *cohesion (cohes)* and *conflict (confl)* subscales, resulting in scores like *fc_p_fes cohes_mean* (mean of items in the parent cohesion subscale) and *fc_p_fes confl_mean* (mean of items in the parent conflict subscale). Each scale/subscale can have one or several summary scores. For example, most means and sums are accompanied by*’nm’* (number missing) scores, which provide the number of items that were missing for the computation of a specific score (e.g., *fc_p_fes cohes_nm*). In the source code, scores are organized by domain, with one R file per domain which are named with the prefix “*scores_*” (e.g., “scores_ab.R”, “scores_fc.R”, *etc*.). There are two exceptions, the “scores_mh_aseba.R” (ASEBA, Achenbach System of Empirically Based Assessment) and “scores_su_sui_tlfb.R” (SUI, Substance Use Interview & Timeline Followback) files, which are separated into their own files due to their complexity and size.

All scoring functions in the package are designed in a consistent way (Fig. S1):

1. Function naming: All functions that compute summary scores begin with the prefix “*compute_*”, followed by the specific score name in the ABCD tabulated dataset. For example, “*compute_mh_p_ders attun_mean*” indicates that it computes the summary score “*mh_p_ders attun_mean*”, the mean of the DERS (Difficulties in Emotion Regulation Scale) attunement subscale.
2. Input: The function expects a data frame containing the variables that are needed for the score calculation. This data frame can be a subset of the larger ABCD dataset, allowing users to focus on specific participants or measurements.
3. Structure of the function: (a) *Validates* the inputs to all parameters, including but not limited to checking if the data frame contains the required variables, if the variables are of the correct type, and if other parameters are within the expected range. If the parameters do not pass validation, the function will return an informative error message to the user. (b) *Computes* the score using the scoring algorithm, and finally, (c) *Returns* the computed score in a desired format.
4. Output: The function returns a data frame containing the computed summary scores, attached to the original data frame (default) or as a separate data frame, depending on the user’s preference.

Functions with the *_all* suffix are high-level convenience functions designed to compute all summary scores within a specific scale/measure at once. For example, “*compute_ph_y_bp_all*” computes all summary scores within the Youth Blood Pressure measure, which includes 6 summary scores: *ph_y_bp sys_mean, ph_y_bp sys_nm, ph_y_bp dia_mean, ph_y_bp dia_nm, ph_y_bp hrate_mean*, and *ph_y_bp hrate_nm* (systolic, diastolic, and heart rate, respectively).

### Scoring Algorithm and Utility Functions

The scoring algorithms were developed in close collaboration with the ABCD Study® domain experts and assessment workgroups. The experts specified the scoring algorithms, handling of missing data, and any specific transformations required. The development team translated these specifications into R code, generated both simulated and real results for the experts to review, and iteratively refined the code based on their feedback. This collaborative approach ensured that the scoring algorithms accurately reflected the intended measures.

To enhance code readability and maintainability, we developed a set of utility functions that are used across multiple scoring algorithms. These utility functions consist of common operations, such as computing means, sums, and the number of missing values, as well as more complex utility functions which, for example, perform row-wise counts of a list of user-defined conditions.

ABCDscores utility functions are designed to be reusable and modular, allowing for easy integration into different scoring algorithms. In ABCDscores, all score-related utility functions are prefixed with “*ss_*” (e.g., *ss_mean, ss_sum, ss_nm, ss_count*). This naming convention clearly indicates their purpose and distinguishes them from other functions in the package. In addition to the summarizing utility functions, ABCDscores contains several utility functions used for data transformations, which provide wrapper functions around powerful tidyverse functions, such as combining two columns into one (*combine_cols*), recoding values in a column (*recode_levels*), and converting a longitudinal variable into a static one (*make_static*). Together, these generic utility functions in ABCDscores can be used to create other summary scores within and beyond ABCD data.

### Documentation

The ABCDscores package is well documented using *roxygen2* comments for function documentation and vignettes providing details about how to use the package. The documentation is also provided as a static website that is created using the *pkgdown* package: https://software.nbdc-datahub.org/ABCDscores/. As part of the function documentation for each score, we provide detailed information about the scoring algorithm, including the rationale behind the scores, and the citations for validated measures that the scores were originally derived from. For scales with more complicated input data requirements and scoring algorithms, e.g., the ASEBA and SUI/TLFB scores, we provide detailed vignettes to guide users through preparing the input data and computing the scores.

### Defaults and Customization

The ABCDscores package is designed to be user-friendly and flexible, allowing users to customize the scoring process according to their specific needs. The package provides default arguments for each scoring function, which are set to recommended values based on experts’ guidelines. These defaults ensure that users can reproduce the exact scores that are published as part of the ABCD tabulated data resource. However, to enable researchers to create scores according to their specific requirements or preferences, we allow users to change these default arguments to suit their specific needs.

All scoring functions in the package have a common set of standard arguments:

- **data**: The input data frame containing the relevant variables for the score calculation. This is a required argument.
- **name**: The name of the score column in the output data frame. The default is always the function name without the “*compute_*” prefix, i.e., the name of the respective score in the ABCD tabulated dataset.
- **combine**: A logical argument indicating whether to combine the computed score with the original data frame. The default is *TRUE*, meaning that the score will be added to the original data frame.

Depending on the specific scoring function, there may be additional default arguments that are specific to the domain or scale, for example, the number of missing items that are allowed to still compute the score. These arguments are described in the function documentation and can be customized as needed.

## Results

### Overview of ABCDscores

As an open-source R package designed to enhance data processing transparency and reproducibility for the ABCD Study® (Fig. 1), ABCDscores calculates summary scores across multiple domains, such as behavioral, cognitive, neuroimaging, and mental health (Table 1). A total of 720 summary score functions and 13 utility functions are included in ABCDscore v6.0.0.

**Table 1.** Number of score functions included in the first release of the package and number of scores computed in the ABCD data release 6.0 by research domain.

| Domain | Functions | Scores |
| --- | --- | --- |
| Neurocognition | 5 | 5 |
| Novel Technologies | 10 | 10 |
| ABCD General | 11 | 11 |
| Physical Health | 90 | 90 |
| Family, Friends & Community | 90 | 90 |
| Mental Health | 454 | 454 |
| Substance Use | 60 | 1030 |
| utility functions | 13 | N/A |
| Total | 733 | 1690 |

**Figure 1.**
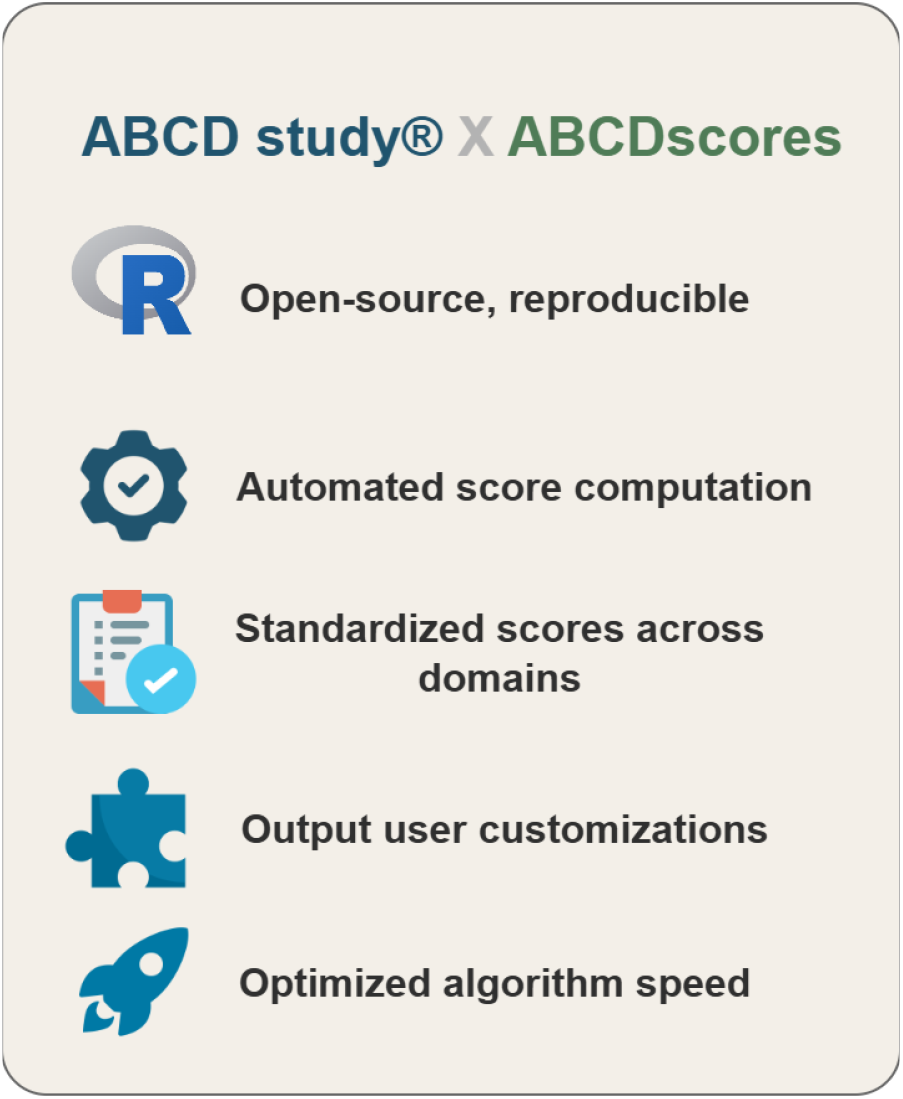
Overview of ABCDscores features in support of the ABCD Study®. This figure highlights the core functionalities of the ABCDscores R package

In most cases, one score function corresponds to one summary score. Exceptions are made in the Substance Use domain. Score functions for two tables, *su_y_sui* (Substance Use Interview [Youth]) and *su_y_tlfb* (Timeline Followback Interview Results [Youth]), can be used to compute multiple summary scores, which causes the number of scores to be much greater than the number of functions in this domain (Table. 1). In total, 1690 scores in the ABCD 6.0 data release were computed using ABCDscores. For future package releases, we expect more functions to be added, empowering numerous new summary scores to be calculated.

Besides the main score functions, the ABCDscores package contains multiple generic, reusable utility functions that will enable users to compute their own summary scores within and beyond the ABCD study®. A list of all generic utility functions available in the ABCDscores package is provided in Table 2. More details about each utility function can be found at: https://software.nbdc-datahub.org/ABCDscores/reference/index.html#utility-functions.

**Table 2.** Overview of utility functions provided in ABCDscores.

| Generic utility functions |  |
| --- | --- |
| Score computation | Data transformation |
| ss_sum | recode_levels |
| ss_mean | combine_cols |
| ss_nm | combine_levels |
| ss_count | make_static |
| ss_count_cond |  |
| ss_max |  |
| ss_prsum |  |
| ss_mean_pos |  |
| ss_tscore |  |

### Performance

Algorithms for calculating the summary scores were optimized to ensure efficient computational time. During the development process, time-intensive steps were identified and substituted with vectorized operations or functions. Consequently, satisfactory processing speed was achieved without the need for multi-core parallelization, allowing the entire dataset (e.g., all scores for the ABCD data release 6.0) to be processed on any modern, everyday-use laptop.

### Use cases

The following examples showcase some basic behaviors and advanced customization of computing summary scores with ABCDscores. In these examples, we use some simulated data for the Family Environment Scale [Parent] cohesion subscale (*fc_p_fes cohes*) to illustrate how most score functions behave and how to apply customizations when using ABCDscores (Fig. 2).

**Figure 2.**
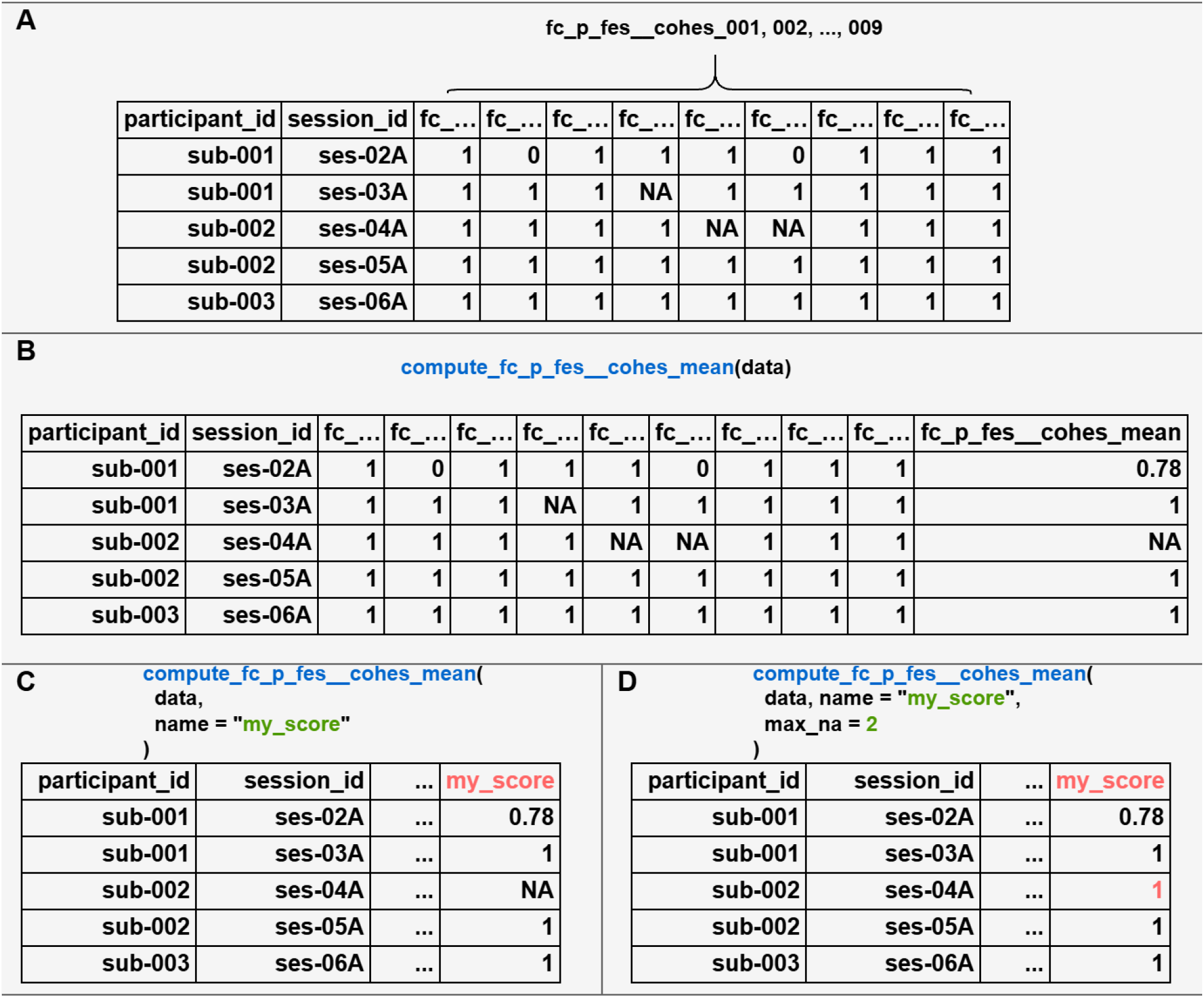
Use case examples of computing scores with the package using simulated data. (A) Input raw data of the Family Environment Scale [Parent] cohesion subscale, which contains 9 items (fc_p_fes cohes_001, 002, …, 009). (B) Example of using all the default arguments to compute the *fc_p_fes cohes_mean* score. (C) Customizing the output column name of the score. (D) Further customizing the maximum number of missing items allowed to 2 (default 1), so some previously *NA* row(s) are calculated with values.

1. To compute the *fc_p_fes cohes_mean* score, the input data frame needs to contain all the required columns. In this case, they are the *fc_p_fes cohes_001, 002, …, 009* columns (Fig. 2A).
2. When the user uses all the default arguments, the only required input for the *compute_fc_p_fes cohes_mean* function is a data frame containing the required columns. The output score is appended to the end of the original input as a new column with name “*fc_p_fes cohes_mean*” (Fig. 2B).
3. As a simple customization, users can, for example, change the column name for the score in the output data frame (Fig. 2C). In this example, the name was changed from “*fc_p_fes cohes_mean*” to “*my_score*”.
4. In the previous example, the final results remained unchanged compared to (2). However, users can also decide to modify parameters that change how the function computes the score and result in different results. In this example, the *max_na* argument (maximum number of missingness allowed) was changed from 1 (default) to 2. As a result, a summary score for the third row is now calculated by the function (Fig. 2D).

In addition to the *max_na* argument, there are several other parameters that can be used to adjust the stringency, range, or variability of scores. All of these parameters come with pre-defined defaults established by the ABCD Study assessment workgroups. Therefore, generating all of the scores from raw data is straightforward, and these advanced parameters are intended for use by experts with particular research objectives.

## Conclusion

ABCDscores is an R package designed to automate the computation of summary scores for the ABCD Study®. By integrating complex scoring algorithms and data processing steps, ABCDscores streamlines analysis workflows, enhances reproducibility, and reduces human error. ABCDscores efficiently handles large datasets, providing accurate and consistent summary scores across various research domains. In conclusion, ABCDscores significantly enhances the analysis of ABCD Study® data, allowing researchers to concentrate on scientific discovery and accelerating insights into adolescent brain and cognitive development.

## Funding

The ABCD Study® Data Analysis, Informatics and Resource Center is supported by the National Institute of Health and additional federal partners under award number: U24DA041123

Additional support for The ABCD Study® is provided by the National Institutes of Health and additional federal partners under award numbers: U01DA041048, U01DA050989, U01DA051016, U01DA041022, U01DA051018, U01DA051037, U01DA050987, U01DA041174, U01DA041106, U01DA041117, U01DA041028, U01DA041134, U01DA050988, U01DA051039, U01DA041156, U01DA041025, U01DA041120, U01DA051038, U01DA041148, U01DA041093, U01DA041089, U24DA041147.

A full list of supporters is available at Federal Partners – ABCD Study® (https://abcdstudy.org/federal-partners.html).

ABCD Consortium investigators designed and implemented the study and/or provided data but did not necessarily participate in the analysis or writing of this report. This manuscript reflects the views of the authors and may not reflect the opinions or views of the NIH or ABCD Consortium investigators.

## Software and data availability

The ABCDscores package can be installed from CRAN and GitHub (https://github.com/nbdc-datahub/ABCDscores/). Documentation and vignettes are available on the package documentation website (https://software.nbdc-datahub.org/ABCDscores/). The package version is associated with the ABCD Study® data release version. For example, the current data release version is 6.0, and the corresponding ABCDscores package version is *6.0.0*. To reproduce scores included in a given data release, use the same version of the package as was used to create the tabulated data for the release. We release minor updates to the package to fix bugs or improve performance. These updates are indicated by the last digit of the version number (e.g., 6.0.1, 6.0.2). Use the most current version of the package to take advantage of the latest fixes and improvements.

## Acknowledgement

We would like to acknowledge the contributions of the ABCD Study® consortium, particularly the assessment workgroups and domain experts who offered valuable input and guidance throughout the development of the ABCDscores package. Additionally, we extend our gratitude to the participants of the ABCD Study® for their time and commitment to advancing science.

## Supplement

**Figure S1.**
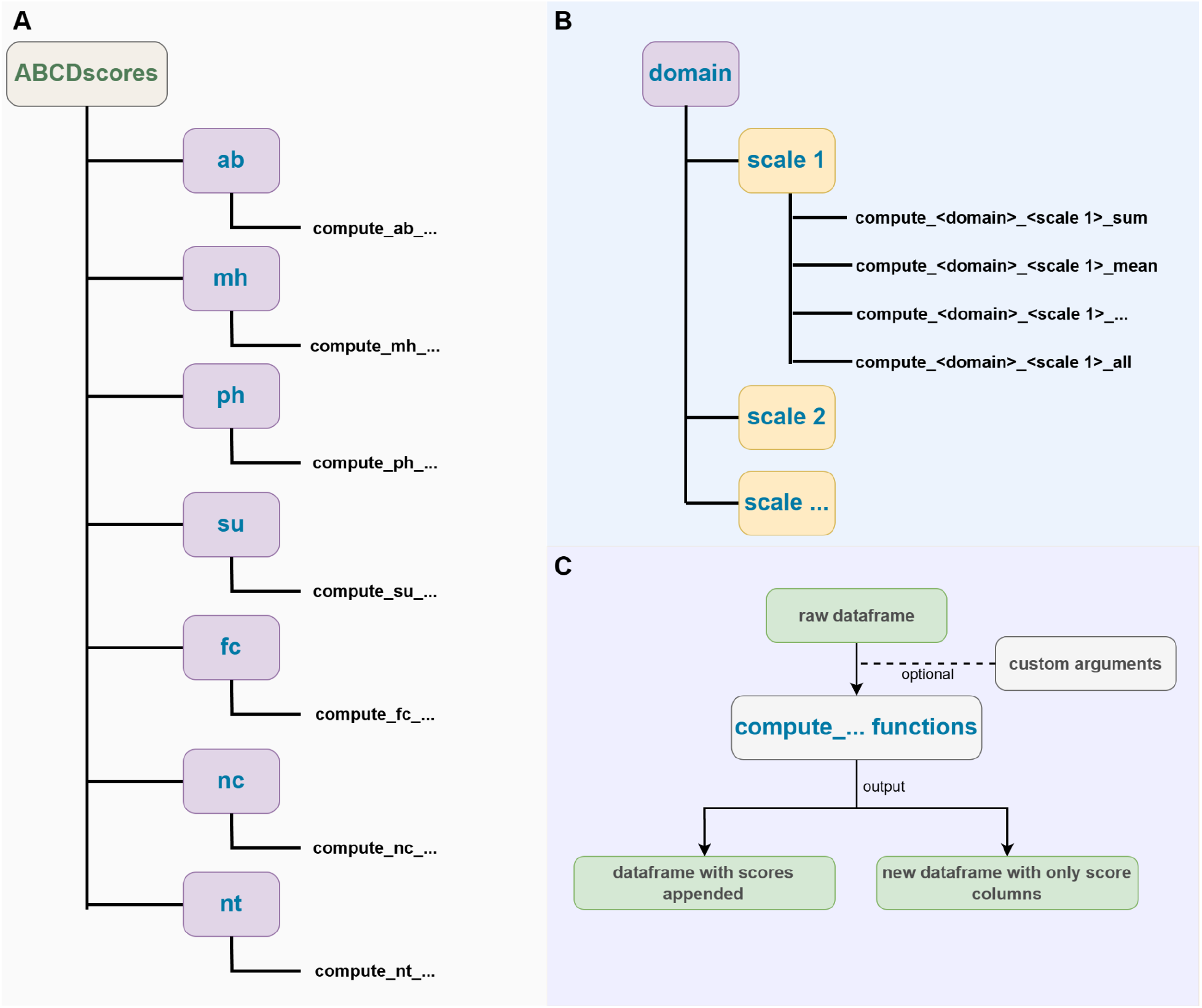
Architecture and workflow of the ABCDscores R package. (A). Overview of the modular structure of ABCDscores. Scores are organized by domain. Each domain (*e.g., ab* for ABCD General, *mh* for Mental Health, etc) includes dedicated functions prefixed with compute_<domain>_… to calculate domain-specific scores. (B). Within each domain, multiple summary score functions, such as sum, mean, and others, are available for each scale or measure. Each function corresponds to an individual score. Additionally, there is an “all” function, indicated by the *“_all”* suffix, which computes all scores within a given scale or measure simultaneously. (C). Scoring workflow: A dataframe containing the required raw score columns is passed to the compute functions, optionally including user-defined arguments. The output can either be the original dataframe with the newly appended score columns or a separate dataframe containing only the calculated scores, depending on the user’s preference.

